# Oral delivery of semaglutide and tirzepatide using milk-derived small extracellular vesicles

**DOI:** 10.1101/2024.12.28.630566

**Authors:** Yuefei Zhang, Jianyi Han, Wei Wu, Bobo Dang

**Affiliations:** Research Center for Industries of the Future and Key Laboratory of Structural Biology of Zhejiang Province, School of Life Sciences, Westlake University; Hangzhou, Zhejiang, China; Westlake Laboratory of Life Sciences and Biomedicine, Hangzhou, Zhejiang, China; Institute of Biology, Westlake Institute for Advanced Study; Hangzhou, Zhejiang, China

## Abstract

Therapeutic proteins and peptides have revolutionized modern biomedicine. However, the large size and complex structure of these macromolecules preclude their oral administration, greatly limiting their applications in patient care. Biochemical degradation in the gastrointestinal tract, mucus, and cellular barrier are the major obstacles to oral development. Small extracellular vesicles (sEVs) are natural nano lipid vesicles serving as essential vehicles for intercellular communication. sEVs are resistant to biochemical degradation, permeable to mucus barriers, and can penetrate cellular barriers. sEVs are thus considered the next-generation vehicles with the potential for oral peptide/protein drug delivery. Herein, we report the use of milk-derived sEVs as delivery vehicles to achieve successful and highly efficient oral delivery of two therapeutic GLP-1 receptor agonists, semaglutide and tirzepatide. We showed that semaglutide and tirzepatide can be efficiently loaded onto sEVs in vitro, and oral gavage of semaglutide-loaded sEVs or tirzepatide-loaded sEVs can effectively lower blood glucose levels in db/db mouse models. This study demonstrates that sEVs is a platform technology for oral peptide drug delivery and opens a new avenue for oral peptide/protein therapeutics delivery.

## Introduction

Since insulin was extracted to treat diabetes, more than 80 peptide drugs have been marketed in the therapeutic areas to treat diabetes, cancer, osteoporosis, etc^1^. However, peptide drugs suffer from very low oral bioavailability, limiting them to invasive injection as the administration route. Parenteral administration of peptide drugs presents significant challenges, leading to reduced drug compliance and limited application^2^. To address this issue, extensive research has been devoted to developing effective oral delivery systems for peptides. This involves utilizing a diverse range of innovative materials and technologies to enhance the oral bioavailability of peptides, focusing on overcoming degradation caused by digestive fluids and enhancing peptide absorption through the gastrointestinal barrier^2^. Despite significant efforts, oral administration of peptides remains a major hurdle, with only a few approved oral peptide drugs exhibiting low oral bioavailability (less than 1%)^3^. The limited oral bioavailability not only compromises the efficacy of these drugs but also necessitates higher dosages for achieving desired therapeutic outcomes^4^.

The first oral GLP-1RA, semaglutide (oral formulation), was approved in 2019 for the treatment of type 2 diabetes, and is considered a breakthrough in the field of oral peptide delivery^3^. The excipient in this formulation, sodium 8-(2-hydroxybenzamido) octanoate (SNAC), effectively increases the local pH around the semaglutide, preventing the pepsin degradation as well as enhancing the absorption through gastric epithelium^2,5^. Despite its advancements, oral semaglutide does exhibit some limitations. The bioavailability of oral semaglutide remains relatively low, ranging from 0.4% to 1%, when compared to its injectable counterpart. This limitation necessitates higher and more frequent dosages, leading to an increased economic burden on patients^6^. The excipient used in the oral formulation fails to adequately enhance the oral absorption of other peptides with structures similar to semaglutide^5^. As a result, this technology cannot be extended to other related peptide drugs, thereby limiting its broader impact and potential applicability beyond semaglutide. In addition, dietary restrictions are required when taking the drug, further adding to the complexities of its use^7^. Moreover, within the realm of glucose-lowering and weight-loss peptide drugs, there is a growing interest in the development of multi-targeted drugs. For instance, tirzepatide, a dual-target agonist of GLP-1R/GIPR, has shown superior glucose-lowering effects than semaglutide in phase III clinical trial^8^. The success of such drugs may likely increase the demand for oral formulations. Consequently, there is a pressing need to develop an efficient, convenient, and universally applicable oral delivery system not only for semaglutide and tirzepatide but also for other related peptides or proteins^3^.

sEVs (small extracellular vesicles) are natural nanovesicles secreted by cells. They bear a similar size and structure to liposomes, but with a more intricate composition, comprising diverse lipids, proteins, and nucleic acids^9^. Functionally, sEVs act as natural carriers of signaling molecules, facilitating cell-to-cell communication in various pathophysiological processes^10^. Being of natural origin, sEVs offer several advantages, including excellent biocompatibility and low immunogenicity when compared to synthetic or viral vehicles. These inherent characteristics position sEVs as promising candidates for the next-generation drug delivery vehicles^11,12^.

Serving as versatile carriers, sEVs are released by almost all cell types, leading to diverse and abundant sEV populations^13^. Additionally, some sEVs have shown remarkable stability in the gastrointestinal tract and the ability to cross biological barriers, rendering them promising candidates for oral drug delivery vehicles^14^. Currently, oral sEVs are primarily sourced from milk, although some studies have explored sEVs derived from cell lines or plants^15,16^. Milk-derived sEVs exhibit low immunogenicity and high stability in the stomach and intestine^17^. The widespread availability and cost-effectiveness of milk provide a strong foundation for the industrial-scale production of milk-derived sEV-based drugs, paving the way for significant advancements in therapeutic applications^12^.

In 2016, a few studies started to use milk sEVs to orally deliver small-molecule chemotherapeutic drugs (including paclitaxel and docetaxel)^18^. Curcumin, anthocyanin^19^, α-mangostin^20^, and resveratrol^21^ were also shown to be orally delivered using milk sEVs. Furthermore, experiments have demonstrated that milk sEVs possess the potential to orally deliver nucleic acid drugs, owing to their resistance to degradation and ability to cross the intestinal barrier^22–27^. In 2022, studies focused on peptide or protein oral delivery by milk sEVs. Oral milk sEVs loaded with insulin were shown to lower blood glucose in type I diabetic rats^28^. Septic mice were relieved by oral FGF21 encapsulated within milk-derived sEVs^29^. Liraglutide was also loaded into milk sEVs for oral delivery, while sublingual delivery was reported to lower blood glucose, oral gavage failed to show significant efficacy^30^. However, among current studies, there is still a lack of comprehensive comparison between different sEV sources or loading methods.

In this study, we investigated the potential for oral delivery of peptide drugs, specifically semaglutide or tirzepatide, utilizing sEVs derived from natural sources. We isolated, purified, and extensively characterized various sEV types obtained from milk (whole or defatted fresh milk), plant (immature or mature coconut water), and the 293F cell line. Among these sources, sEVs from defatted milk exhibited notably high yields and demonstrated favorable characteristics in terms of particle size, morphology, marker expression, and purity. As a result, we selected defatted milk-derived sEVs as the optimal oral delivery vehicles for our experimental peptide drugs. Through a thorough comparison of various loading methods, we discovered that room temperature (RT) incubation produced sEV-semaglutide complex with the highest encapsulation rate (56.4%) and high positive rate (94.9%). Both in vitro and in vivo assays demonstrated that sEV-semaglutide retained a similar potency to standalone semaglutide. We obtained similar results in sEV-tirzepatide and tirzepatide. Moreover, in vivo distribution analysis revealed that the capsule sEV formulation persisted significantly longer than the liquid form after oral administration. Furthermore, an in vivo efficacy study in a diabetic mouse model showcased the robust performance of the oral capsule formulations containing sEV-semaglutide and sEV-tirzepatide. These formulations demonstrated significant and sustained blood glucose-lowering effects. The compelling results indicate that the milk sEV capsules we developed have the potential to enhance the oral absorption and efficacy of both semaglutide and tirzepatide, offering promising prospects for advanced oral peptide drug delivery.

## Results

### Defatted milk sEVs are utilized as oral delivery vehicles

Currently, the sources of oral sEVs are mainly focused on milk, with a few derived from cell lines or plants^15,16^. However, these sEVs have not been studied comparatively. In this study, we focused on representative sEVs from milk (whole or defatted), coconut water (immature or mature), and 293F cells^31–33^. A comprehensive analysis was conducted through nanoparticle tracking analysis (NTA), transmission electron microscopy (TEM), and western blot. We analyzed the particle sizes of the sEVs, which were shown to be around 100-200 nm (Figure 1A). TEM data showed that the sEVs we isolated have a “cup-shaped” structure (Figure 1B). We found that milk sEVs have the highest yields: the yields of defatted milk sEV reach 10^11^ sEVs/mL, which is slightly higher than whole milk sEVs (Figure 1A). We thus selected milk sEVs for further characterizations. Phospholipid bilayers of milk sEVs can be clearly observed by cryo-EM (Figure 1C). TEM and cryo-EM show that there are more lipoprotein particles (indicated by black arrowheads) in whole milk sEV samples compared with sEVs from defatted milk, suggesting low purity of whole milk derived sEVs (Figure 1B-C). We then analyzed the expression of generally recommended sEV-positive markers (CD9, TSG101) and negative marker (GM130). The results showed that the expression of positive marker proteins in defatted milk sEVs was higher than that in whole milk derived sEVs (Figure 1D and Figure S1). The data indicated that defatted milk sEVs exhibited higher purity compared to whole milk sEVs. Considering factors such as yield, morphology, and purity, we selected defatted milk-derived sEVs as the preferred oral delivery vehicles. Henceforth, sEVs will refer specifically to defatted milk sEVs.

**Figure 1.**
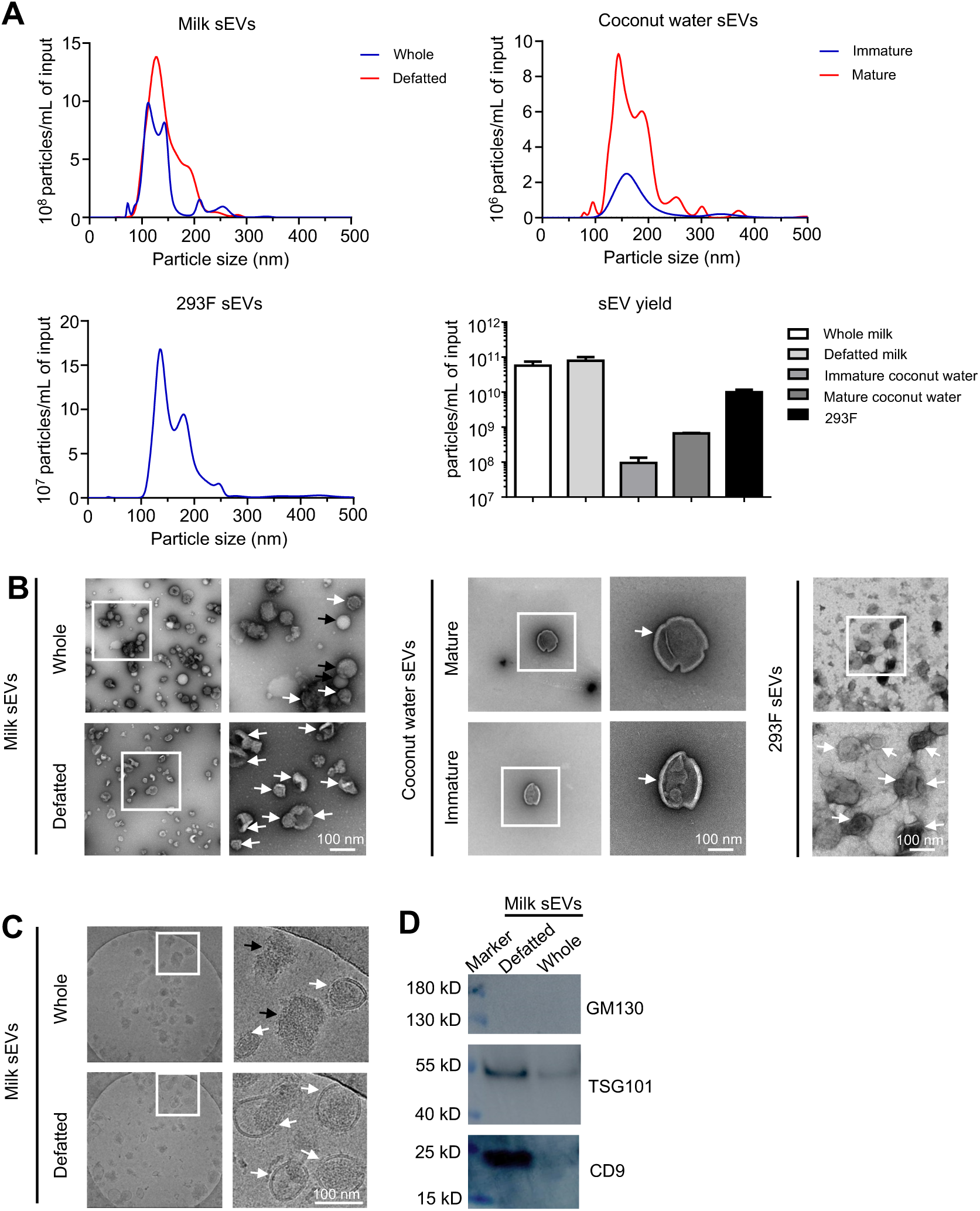
sEV isolation and characterization. (A) sEV size distribution and yield determined by nanoparticle tracking analysis (n = 3 to 7 per group). Morphological characterization of sEVs through TEM (B) or cryo-EM (C). White arrows indicate sEVs with cup-shape in TEM or lipid bilayer in cryo-EM. Black arrows indicate lipoprotein particles. (D) Western blot analysis of sEV positive marker CD9, TSG101 and negative marker GM130 (35 μg total protein loaded per sample).

### In vitro loading of semaglutide or tirzepatide into sEVs

The methods used to load drugs into sEVs include incubation, sonication, extrusion, freeze-thaw, and detergent treatment^34^. We used encapsulation efficiency and positive rate to compare different loading methods. The encapsulation efficiency is defined by the concentration of the sEV-incorporated drugs over the initial concentration used to prepare the formulation^35^. The positive rate represents the ratio of drug-loaded sEVs to the total sEVs. To track and quantify semaglutide or tirzepatide, we synthesized semaglutide-FAM (Figure S2) or tirzepatide-FAM (Figure S3) to monitor the loading process. sEVs are much larger than semaglutide or tirzepatide in size. So, the elution peak of sEVs on fast protein liquid chromatography (FPLC) is before 10 mL, while semaglutide or tirzepatide is eluted after 20 mL (Figure S4). Therefore, we can calculate the encapsulation efficiency by determining the relative proportions of FAM intensity at the above two positions. We loaded semaglutide-FAM into sEV using incubation, sonication, extrusion, freeze-thaw, and saponin treatment methods, and calculated encapsulation efficiency. The results showed that incubation at room temperature has the highest encapsulation efficiency of 56.4% (Figure 2A). The sEV-semaglutide-FAM has a positive rate of 94.9% determined by nanoFCM (Figure 2B). We also observed colocalization of semaglutide-FAM and sEVs (Figure 2C) using super-resolution imaging. In addition, sEV-tirzepatide-FAM prepared by incubation has 23% encapsulation efficiency and 83.4% positive rate, respectively (Figure S5). The above data indicated that we can successfully prepare sEV-semaglutide or sEV-tirzepatide by incubation at room temperature.

**Figure 2.**
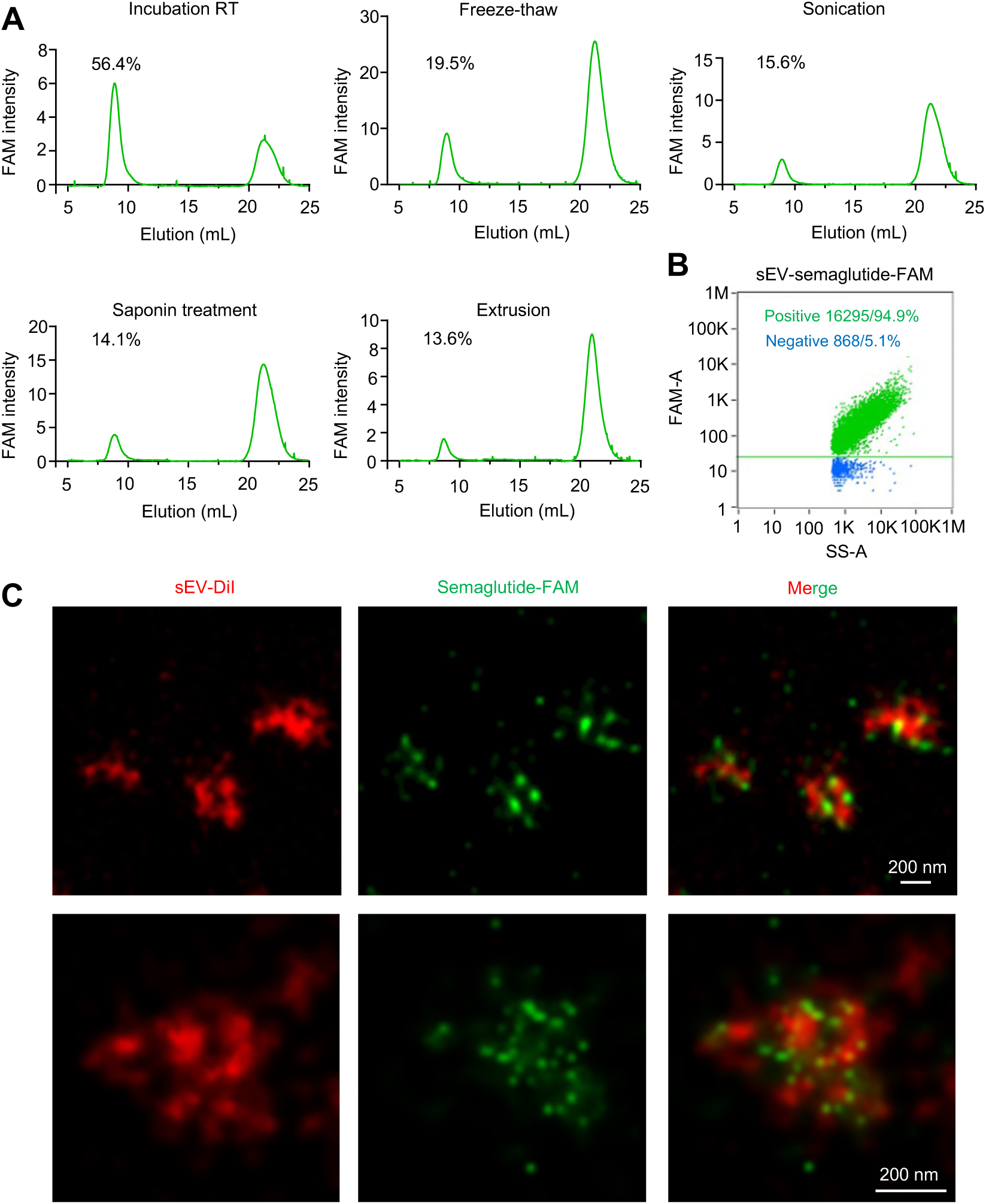
Production of sEV-semaglutide. (A) Efficiency of loading semaglutide-FAM into sEVs. (B) Positive rate of sEVs loaded with semaglutide-FAM. (C) Super-resolution microscopy of sEVs loaded with semaglutide-FAM.

### In vitro and in vivo activity of sEV-semaglutide and sEV-tirzepatide

To investigate the in vitro and in vivo activity of sEV-semaglutide or sEV-tirzepatide, we utilized an in vitro cell assay (293 cell line that expresses both the hGLP-1R and CRE firefly luciferase) and an in vivo mouse model (db/db diabetic mice)^36^. We mixed semaglutide with semaglutide-FAM in proportion (10:1) and then incubated with sEVs at room temperature. The fluorescent standard curve of semaglutide-FAM allowed us to calculate the amount of semaglutide in the sEV-semaglutide samples.

Cell assays revealed that sEV-semaglutide and sEV-tirzepatide exhibited EC_50_ values comparable to those of the peptide-only groups (Figure 3A-B). Furthermore, subcutaneous injection results showed that sEV-semaglutide exhibited glucose-lowering effects similar to semaglutide (Figure 3C). Similarly, in the tirzepatide group, sEV-tirzepatide effectively reduced blood glucose levels (Figure 3D). These findings indicate that sEV-peptides possess comparable efficacy to their peptide-only counterparts.

**Figure 3.**
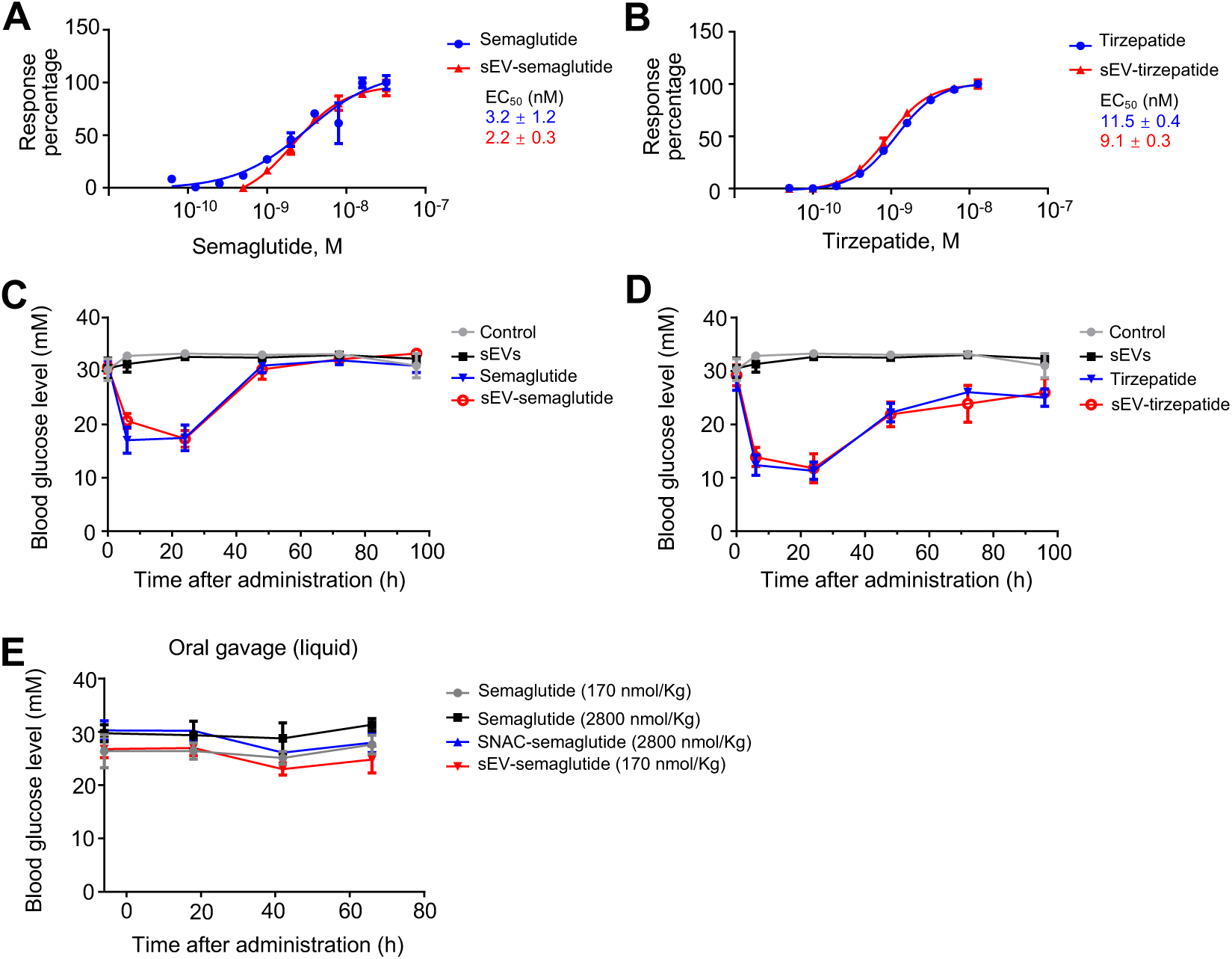
Efficacy of sEV-peptides. (A) GLP-1R activation (EC_50_) of semaglutide and sEV-semaglutide. (B) GLP-1R activation (EC_50_) of tirzepatide and sEV-tirzepatide. (C) Blood glucose lowering efficacy study of semaglutide and sEV-semaglutide in diabetic db/db mice after s.c. dosing (30 nmol/Kg, n =5 to 6 per group). (D) Blood glucose lowering efficacy study of tirzepatide and sEV-tirzepatide in diabetic db/db mice after s.c. dosing (30 nmol/Kg,n =5 to 6 per group). (E) Blood glucose level of diabetic db/db mice after p.o. dosing with semaglutide or sEV-semaglutide in liquid formulation, supplemented with SNAC or nothing (n =4 to 6 per group).

The approved oral semaglutide (Rybelsus) is administered at a dose of 5 mg per individual, equivalent to approximately 17 nmol/kg for a 70 kg body weight^5^. In a diabetic mouse model, we initially attempted oral delivery of a 10-fold dose (170 nmol/kg) of semaglutide or sEV-semaglutide in a liquid formulation. However, neither group exhibited a significant reduction in blood glucose levels (Figure 3E). SNAC is the critical enhancer in promoting oral absorption of semaglutide^5^. We next tested an increased dose of oral semaglutide (2800 nmol/kg) in a solution supplemented with SNAC. However, this formulation still failed to lower glucose levels compared to the control group. In the commercial oral formulation, semaglutide is co-formulated with SNAC in tablet form. Mechanistic studies have shown that semaglutide absorption occurs in the stomach, where the oral semaglutide tablets remain and sustain a high local drug concentration^5^. We hypothesize that the solid formulation prolongs the residence time of semaglutide in the digestive tract, facilitating its absorption. Consequently, the failure to reduce blood glucose levels in our study was likely attributable to the use of the liquid formulation.

### sEV capsules have longer half-life than the liquid formulation in vivo

We then lyophilized semaglutide-loaded sEVs into a dry powder for oral capsule preparation. Direct lyophilization led to a reduction in sEV concentration and altered its size distribution. However, these adverse effects were mitigated by the addition of the cryoprotectant trehalose (TRE) (Figure 4A, Figure S6).

**Figure 4.**
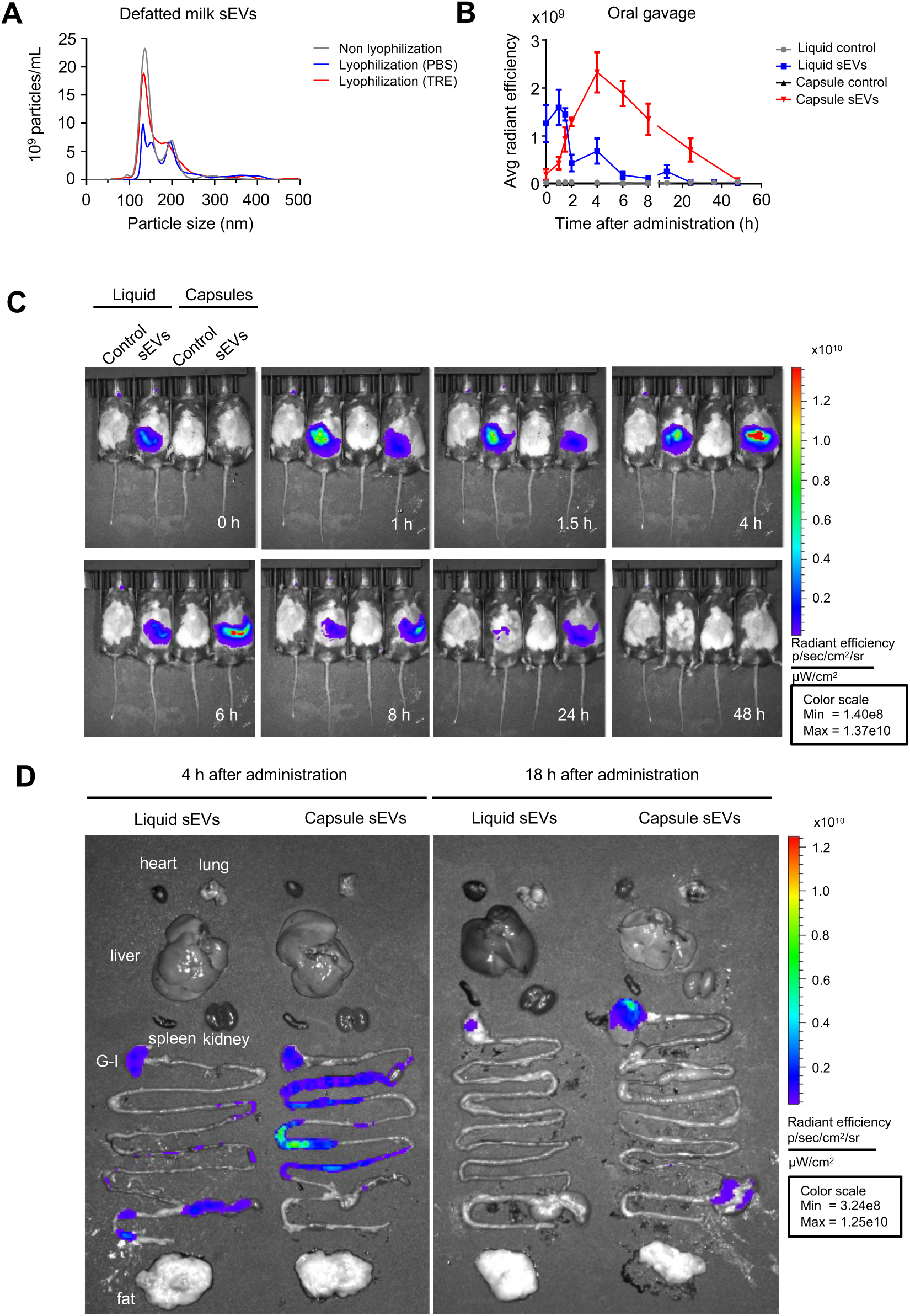
Biodistribution of orally administered bovine milk-derived sEVs. (A) Analysis of sEV size distribution and concentration, both with and without lyophilization, was performed to assess the impact of the lyophilization process. (B-C) db/db mice were administered a single dose of VivoTrack680-labeled sEVs (≈3×10^11^ particles) by oral gavage (p.o.) in liquid or capsule formulation. In vivo imaging and statistical analysis of the mice after drug administration (n = 3 to 5 per group). (D) Representative ex vivo imaging of the tissues and organs.

Additionally, we prepared VivoTrack680-labeled sEVs to investigate the oral absorption process of both sEV liquid and capsule formulations^37^. The signal from the liquid formulation rapidly diminished following oral administration, with most of the signal disappearing within 8 hours, indicating that a significant amount of sEVs were excreted from the body. In contrast, the sEV signal from the capsule formulation showed an initial increase followed by a decrease, with the peak signal observed around 4 hours (Figure 4B). The signal from the capsule formulation remained stronger than that of the liquid formulation from 2 to 48 hours (Figure 4C). Organ distribution analysis following oral administration revealed an enhanced signal in the gastrointestinal tract for the capsule group (Figure 4D). These results suggest that capsule-formulated sEVs persist longer in vivo compared to the liquid formulation.

### sEV-semaglutide and sEV-tirzepatide oral capsules lower blood glucose level

We then repeated the efficacy study using capsulated sEV-semaglutide and sEV-tirzepatide powders, comparing them with semaglutide and SNAC-semaglutide in db/db mouse models. The oral enhancer SNAC was critical for the successful oral delivery of semaglutide, as evidenced by the expected blood glucose-lowering effect in the SNAC-semaglutide group, while the control group without SNAC showed no effect. Both sEV-semaglutide and sEV-tirzepatide significantly reduced blood glucose levels compared to the control group (Figure 5), suggesting that sEVs may play a similar role as SNAC in enhancing the oral availability of semaglutide, thereby reducing blood glucose levels. Moreover, sEV-tirzepatide exhibited a more profound and long-lasting effect in this diabetic mouse model, as evidenced by the lower blood glucose levels at later time points (Figure 5). In conclusion, milk-derived sEVs enhanced the oral delivery of semaglutide and tirzepatide, demonstrating significant potential for the development of oral peptide therapeutics.

**Figure 5.**
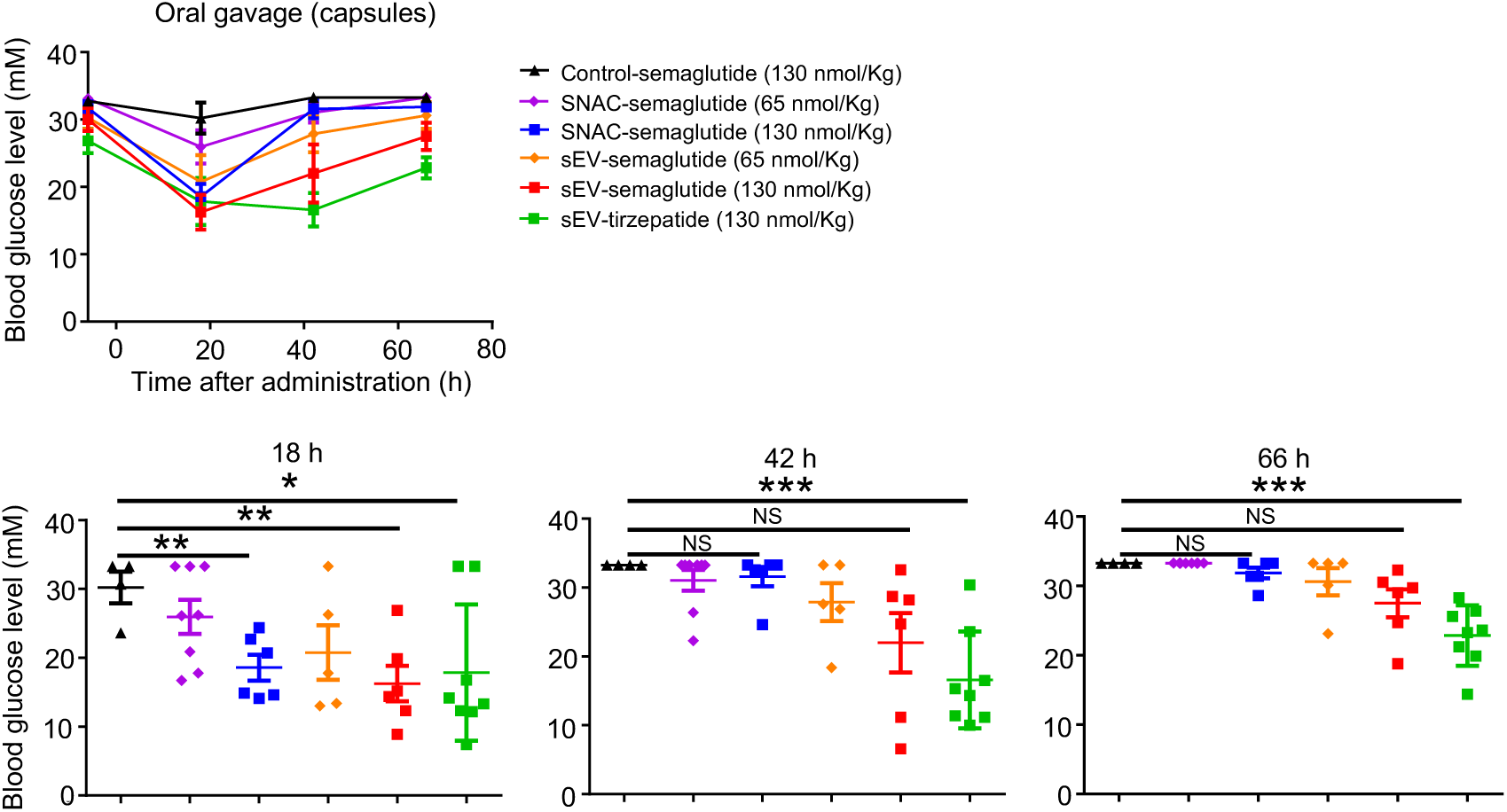
Oral capsules containing sEV-semaglutide and sEV-tirzepatide effectively lowered blood glucose levels in db/db mouse models. Different formulations of semaglutide and tirzepatide were administered orally to db/db mice (n = 4-8). Blood glucose levels were monitored at various time points and compared between different groups.

## Discussion

Peptide drugs have long been difficult to be utilized orally and only a few oral formulations have been approved^3^. In this study, sEV capsules were prepared to achieve oral delivery of semaglutide and tirzepatide, lowering blood glucose level in a diabetic mouse model.

Liraglutide, a GLP-1 receptor agonist, has also been tested in milk sEV-liraglutide oral liquid formulations, but it failed to lower blood glucose levels in mouse models, consistent with our own finding that liquid formulations are ineffective for oral delivery^30^. Previous studies have shown that SNAC can promote the oral delivery of semaglutide but not liraglutide, highlighting the limited applicability of SNAC for oral delivery beyond semaglutide^5^. Recently, milk exosome-liposome hybrid vesicles have been explored for the oral delivery of semaglutide, but their broader application remains uncertain, with the complexity of the delivery system posing additional challenges for future development^38^. In comparison, the sEV capsules we developed not only facilitate the oral delivery of semaglutide but also of tirzepatide, suggesting that sEV capsule formulations hold greater potential for the oral delivery of related peptide drugs^39^.

Extraction methods influence the yield or oral performance of milk sEVs^40^. So, it should be noted that relevant changes likely will improve the oral delivery of milk sEVs. Because of heterogeneity and subpopulations^41,42^, it is worth exploring the subtypes of milk sEVs suitable for oral delivery. In addition, more formulations can be studied in sEV-mediated peptide oral delivery, such as layer-by-layer self-assembly or hydrogel^43,44^.

Overall, we conducted comprehensive studies of sEV sources, loading methods, parameter characterization, in vivo distribution, and formulation studies in developing oral sEV-semaglutide or sEV-tirzepatide. These oral capsule formulations can successfully reduce blood glucose level in mouse model, which has the potential to be a practical solution for oral delivery of peptide or protein payloads.

## Acknowledgments

We thank the Mass Spectrometry & Metabolomics Core Facility, Laboratory Animal Resources Center and Center for Biomedical Research Core Facilities of Westlake University for sample analysis.

## Author contributions

B.D., Y.Z. designed the project. B.D., Y.Z. analyzed the data, wrote, and revised the manuscript.

